# Comparative analysis of seven short-reads sequencing platforms using the Korean Reference Genome: MGI and Illumina sequencing benchmark for whole-genome sequencing

**DOI:** 10.1101/2020.03.22.002840

**Authors:** Hak-Min Kim, Sungwon Jeon, Oksung Chung, Je Hoon Jun, Hui-Su Kim, Asta Blazyte, Hwang-Yeol Lee, Youngseok Yu, Yun Sung Cho, Dan M. Bolser, Jong Bhak

**Author notes:** Email address: H.M.K., S.J., O.C., J.H.J., H.S.K., A.B., H.Y.L., Y.Y., Y.S.C., D.M.B., J.B.

## Abstract

**Background:** MGISEQ-T7 is a new whole-genome sequencer developed by Complete Genomics and MGI utilizing DNA nanoball and combinatorial probe anchor synthesis technologies for generating short reads at a very large scale – up to 60 human genomes per day. However, it has not been objectively and systematically compared against Illumina short-read sequencers.

**Findings:** By using the same KOREF sample, the Korean Reference Genome, we have compared seven sequencing platforms including BGISEQ-500, MGISEQ-T7, HiSeq2000, HiSeq2500, HiSeq4000, HiSeqX10, and NovaSeq6000. We measured sequencing quality by comparing sequencing statistics (base quality, duplication rate, and random error rate), mapping statistics (mapping rate, depth distribution, and %GC coverage), and variant statistics (transition/transversion ratio, dbSNP annotation rate, and concordance rate with SNP genotyping chip) across the seven sequencing platforms. We found that MGI platforms showed a higher concordance rate of SNP genotyping than HiSeq2000 and HiSeq4000. The similarity matrix of variant calls confirmed that the two MGI platforms have the most similar characteristics to the HiSeq2500 platform.

**Conclusions:** Overall, MGI and Illumina sequencing platforms showed comparable levels of sequencing quality, uniformity of coverage, %GC coverage, and variant accuracy, thus we conclude that the MGI platforms can be used for a wide range of genomics research fields at approximately half the cost of the Illumina platforms.

## Introduction

Recently, due to the rapid technological advancement, the second- and third-generation sequencing platforms can produce a large amount of short- or long-reads data at relatively low cost [1]. Depending on the application, these sequencers offer several distinct advantages. Short-read based second-generation sequencing can be used to efficiently and accurately identify genomic variations. Long-read based third-generation sequencing can be used to identify structural variations and build high quality *de novo* genome assemblies [2]. Shortread sequencing technologies are routinely used in large-scale population analyses and molecular diagnostic applications because of the low cost and high accuracy [3]. The most commonly used platforms from Illumina are the HiSeqX10 and NovaSeq6000 short-read sequencers. A competing sequencer developed by Complete Genomics and MGI Tech is the MGISEQ-T7. MGISEQ-T7 is a new sequencing platform after BGISEQ-500 that uses DNA nanoball and combinatorial probe anchor synthesis to generate short reads at a very large scale [4]. In the present study, we compared seven short-read based sequencers; two MGI platforms (BGISEQ-500 and MGISEQ-T7) and five Illumina platforms (HiSeq2000, HiSeq2500, HiSeq4000, HiSeqX10, and NovaSeq6000) (Table 1), in terms of their base quality, uniformity of coverage, %GC coverage, and identification of the variants.

**Table 1.**
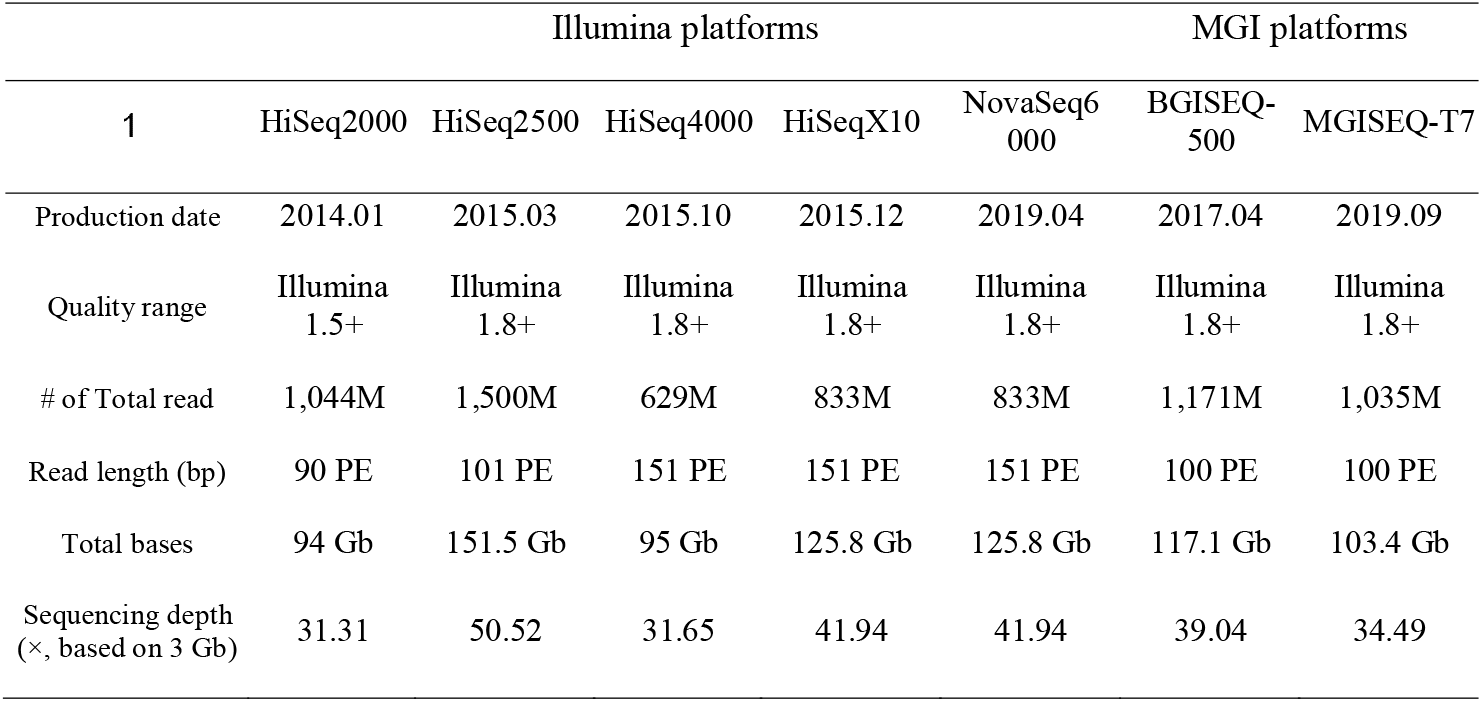
Raw read statistics for seven sequencing platforms

## Results

### Sequencing data summary

We analyzed and benchmarked the whole-genome sequencing (WGS) data quality generated by the seven sequencers using the KOREF (the Korean Reference Genome) [5] DNA. Due to the sequential release and distribution of the sequencers, KOREF sequencing has been carried out in nine years since 2010. Therefore, the blood samples, library construction, and sequencing conditions were not the same. The Illumina platform data used here were from 2014 to 2019, while the MGI platform data were from 2017 and 2019. Also, the read length differs depending on the platform. The Illumina HiSeq2000 had the shortest read length of 90 bp paired-end (PE) and the HiSeq4000, HiSeqX10, and NovaSeq6000 had 151 bp PE. The read length of the HiSeq2500 is 101 bp PE and that of the BGISEQ-500 and MGISEQ-T7 is 100 bp PE. Also, there is a difference in the amount of data as well. Thus, we randomly selected 35× coverage sequencing data for HiSeq2500 and NovaSeq6000 which did have that much sequencing data. HiSeq2000, HiSeq4000, and MGISEQ-T7 had roughly 30× coverage.

### Assessment of base quality and sequencing error of raw reads

Base quality is an important factor in evaluating the performance of sequencing platforms. We analyzed the sequencing quality by identifying the low-quality reads. First, we investigated the base quality distribution of raw reads with the FastQC (FastQC, RRID:SCR_014583) [6]. All the seven platforms showed that the quality of each nucleotide gradually decreased towards the end of a read (Fig. S1). The quality value of the HiSeq4000 and HiSeqX10 reads showed a tendency to decrease rapidly at the end of the read. We defined low-quality reads as those that had more than 30% of bases with a sequencing quality score lower than 20. The fraction of low-quality reads ranged from 2.8% to 18.3% across the seven platforms (Fig. S2 and Table S1). Based on the filtering criteria, the newest platforms, NovaSeq6000 and MGISEQ-T7, showed the lowest percentage of low-quality reads (2.8% and 4.2%, respectively).

We analyzed the frequency of random sequencing errors (ambiguous base, N), which is also an important factor to evaluate the quality of the sequencing platform. We found that the HiSeq2000, HiSeq4000, and HiSeqX10 showed a high random error ratio in certain sequencing cycles (Fig. S3 and Table S2). Furthermore, in the case of HiSeq2000, the random error tended to increase gradually after each sequencing cycle. We also investigated the sequencing error by *K*-mer analysis. Most erroneous *K*-mers caused by sequencing error appear in very low frequency and form the left-side sharp peak [7, 8]. Distribution of *K*-mer frequency showed similar distributions between the platforms (Fig. 1). However, there was a difference in the proportion of low-frequency *K*-mer (≤ 3 *K*-mer depth), which was considered as putative sequencing errors (Table S3). The NovaSeq6000 showed the lowest amount of erroneous *K*-mer (3.91%), while the HiSeq4000 contained the highest amount of erroneous *K*-mer (13.91%) among the seven platforms. The BGISEQ-500 and MGISEQ-T7 showed a moderate level of erroneous *K*-mer (7.72% and 6.39%, respectively).

**Figure 1.**
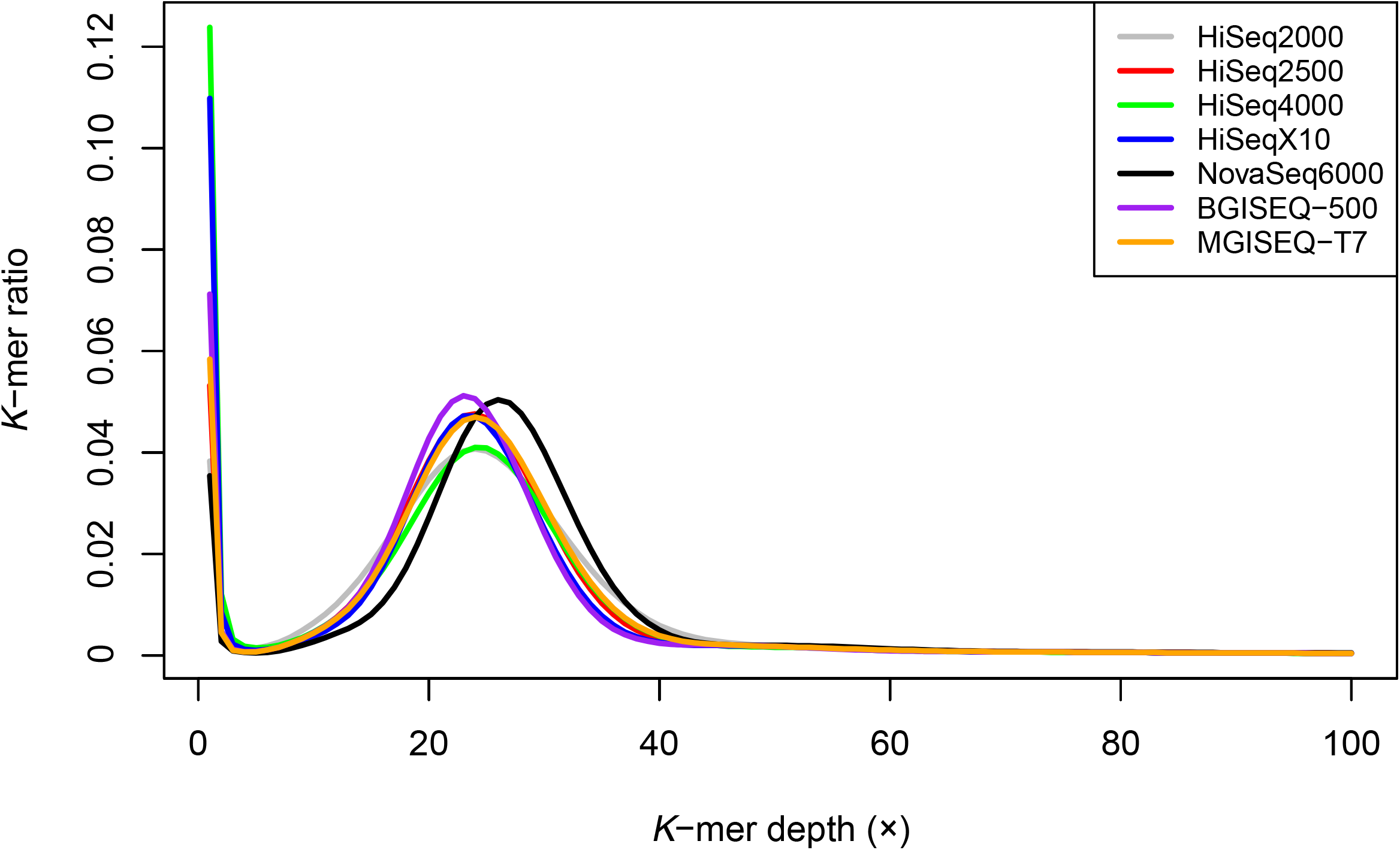
Distribution of *K*-mer frequency for 21-mers using raw reads from seven sequencing platforms. The x-axis represents *K*-mer depth, and the y-axis represents the proportion of *K*-mer, as calculated by the frequency at that depth divided by the total frequency at all depths.

We examined the PCR duplication and adapter contamination in the seven sequencing platforms (Table S2). The HiSeq2000 and MGISEQ-T7 showed the highest duplicate ratio (8.71% in HiSeq2000 and 3.04% in MGISEQ-T7). The HiSeq4000, HiSeqX10 and NovaSeq6000 showed higher adapter contamination rates than other platforms, probably due to longer sequence length (151 bp). However, duplicates and adapter contamination may be more affected by the process of sample preparation than by the sequencing instrument.

### Genome coverage and sequencing uniformity

In order to further assess the genomic coverage and sequencing uniformity, we aligned quality-filtered reads to the human reference genome (GRCh38). Prior to aligning clean reads to the human reference genome, the clean reads of HiSeq2500 and NovaSeq6000 were downsampled to 35 × depth for a fair comparison with the other platforms (Table S4). All seven platforms showed a mapping rate of more than 99.98% and genome coverage of more than 99.6% (≥ 1×; Table 2). We observed a higher duplicate mapping rate in the HiSeq2000 (15.35%) and MGISEQ-T7 (8.77%) than the other platforms and the same pattern as the duplication rates of raw reads (see Table S2). The insert-size for paired-end libraries corresponds to the targeted fragment size for each platform (Fig. S4). It is reported that the depth of coverage is often far from evenly distributed across the sequenced genome [9]. To assess the sequencing uniformity, we analyzed the distribution of mapping depth for all chromosomes (Fig. S5). All seven platforms showed a similar pattern of depth distribution, but interestingly, we found that the depth near the centromere regions was lower exclusively in the HiSeq4000 (Figs. S6-S9). We speculate that this may have been due to a bias in the library preparation step on the HiSeq4000 platform.

**Table 2.**
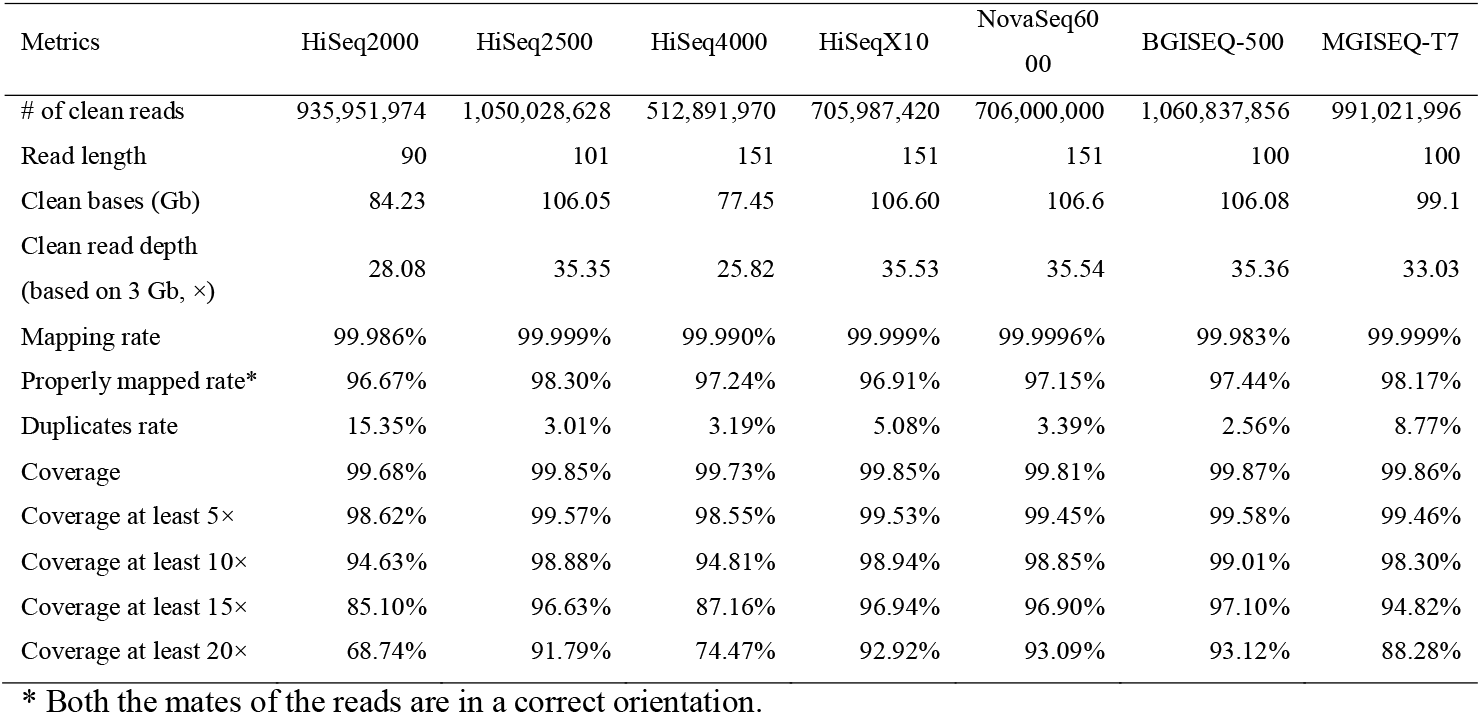
Mapping and coverage statistics

In order to examine the platform-specific covered region of MGI and Illumina platforms, we defined a platform-specific covered region that had significantly different depths (five times difference with an average depth between MGI and Illumina platforms) based on the 100 bp non-overlapping windows. We found 151 Kb and 226 Kb of the platform-specific covered regions from MGI and Illumina platforms, respectively (Table S5). A total of 243 and 717 genes were overlapped in MGI and Illumina specific covered regions, respectively, and most of them were intronic. However, interestingly, the platform-specific covered regions showed a significantly different distribution of GC ratios between the MGI and Illumina platforms (Fig. S10). The MGI platforms tend to cover regions relatively high in GC content (Wilcoxon rank-sum test, *P* = 7.06 × 10^-143^). Nevertheless, it is obvious that platform-specific covered regions for Illumina platforms are slightly longer than those of the MGI platforms, and these regions were not sufficiently covered by the MGI platforms.

Biases in PCR amplification create uneven genomic representation in classical Illumina libraries [10, 11] as PCR is sensitive to extreme GC-content variation [12]. Thus, we analyzed the GC biases in seven platforms. We examined the distribution of GC content in sequencing reads and found that raw reads of all seven sequencing platforms showed a similar GC content distribution to the human reference genome (Fig. S11). To better understand what parts of the genome were not covered properly, we generated GC-bias plots, showing relative coverage at each GC level. Unbiased sequencing would not be affected by GC composition, resulting in a flat line along with relative coverage = 1. We found that all seven platforms provided nearly even coverage at the moderate-GC range 20% to 60%, which represents approximately 95% of the human genome (Fig. 2). On the other hand, the relative coverage of the HiSeq2000 platform dropped more dramatically above 60% GC than other platforms, while the NovaSeq6000 covered well above 60% GC, unlike the other platforms.

**Figure 2.**
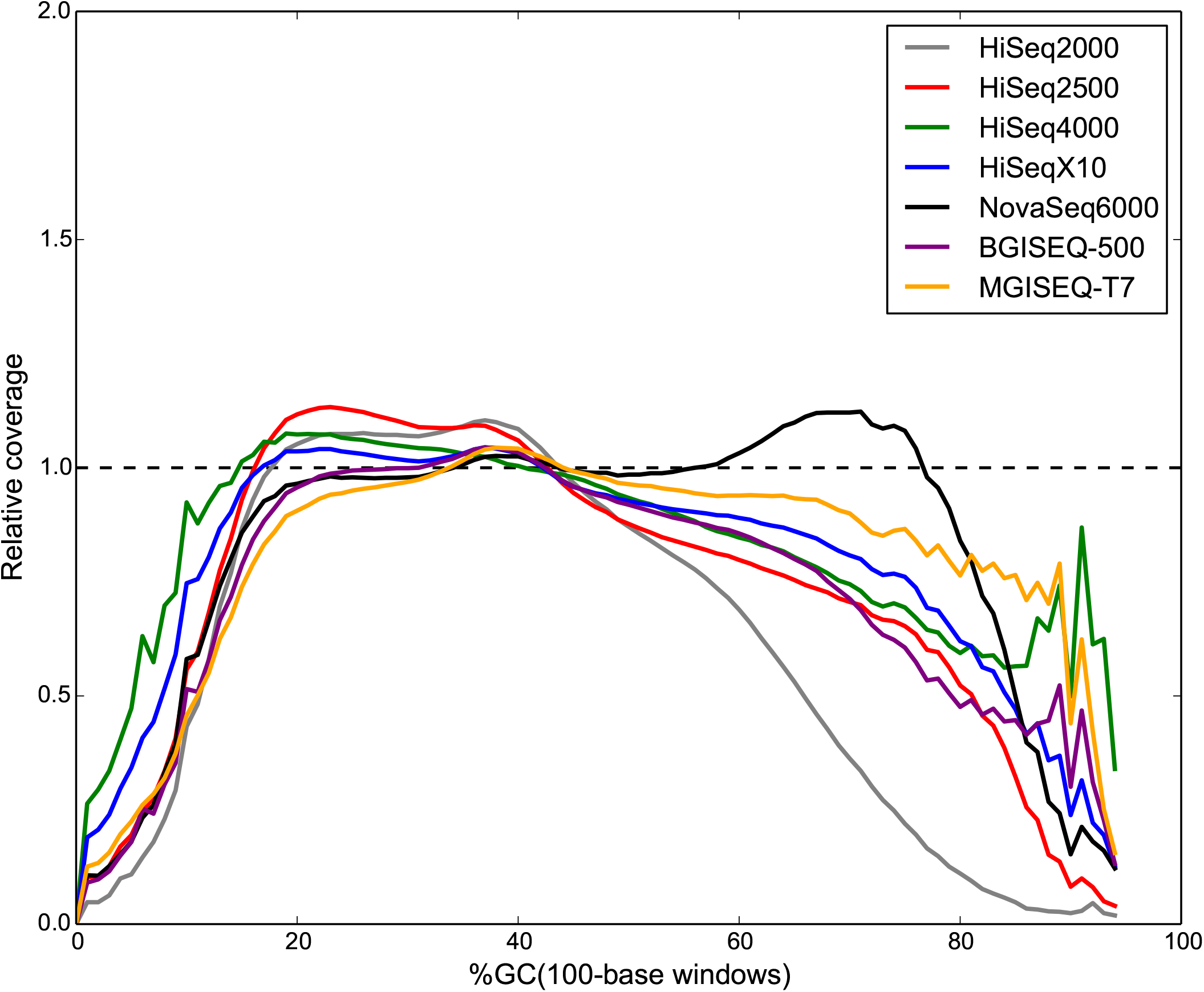
GC-bias plots for seven sequencing platforms. Unbiased coverage is represented by a horizontal dashed line at relative coverage = 1. A relative coverage below 1 indicates lower than expected coverage and above 1 indicates higher than expected coverage.

### Comparison of variants detected among seven platforms

To investigate the performance of variant calling in seven platforms, we adapted the widely used pipeline BWA-MEM (BWA, RRID:SCR_010910) [13] and GATK (GATK, RRID:SCR_001876) [14–16]. We identified an average of 4.18 million single nucleotide variants (SNVs), and 0.66 million indels (insertion and deletion) on each of the seven platforms (Table 3). The statistics of SNVs were similar across all seven in terms of the dbSNP annotation rate (dbSNP153) and the transition/transversion (Ti/Tv) ratio, which indirectly reflects SNV calling accuracy. About 3.7 million SNV loci were found on all seven platforms and this accounts for 87% to 91% of the discovered SNVs on each platform (Table S6). We found 15,670 and 9,325 platform-specific SNVs on the MGI and Illumina platforms, respectively. Interestingly, the number of singletons, variations found only on one platform, was higher for the Illumina (~0.10 million SNVs on average) than MGI (~0.05 million SNVs on average; Table S7). This means that the difference within the Illumina platforms is greater than the difference between the MGI platforms. We also analyzed the number of SNVs found in any six of the seven platforms, which we considered as false negatives. The HiSeq2000 had the largest number of false negatives (79,982 SNVs) among the seven platforms. The two MGI platforms (MGISEQ-T7 and BGISEQ-500) had 16,328 and 10,595 false negatives, respectively, and those of the NovaSeq6000 showed the smallest number of false negatives (4,237 SNVs). To investigate the relationship between the sequencing platforms, an unrooted tree was constructed using a total of 1,034,447 loci where the genotypes of one or more platforms differ from the rest of the platforms (Fig. 3 and Table S8). We found that the two MGI platforms grouped together and they are the closest to the Illumina HiSeq2500 platform. The Illumina platforms were divided into two subgroups in the tree: a long-read length (151 bp) group, containing the HiSeq4000, HiSeqX10, and NovaSeq6000 platforms and a shortread length (<101 bp) group, containing the HiSeq2000 and HiSeq2500 platforms. Read length primarily affects the detection of variants through alignment bias and alignment errors, which are higher for short reads because there is less chance of a unique alignment to the reference sequence than with longer reads [17].

**Figure 3.**
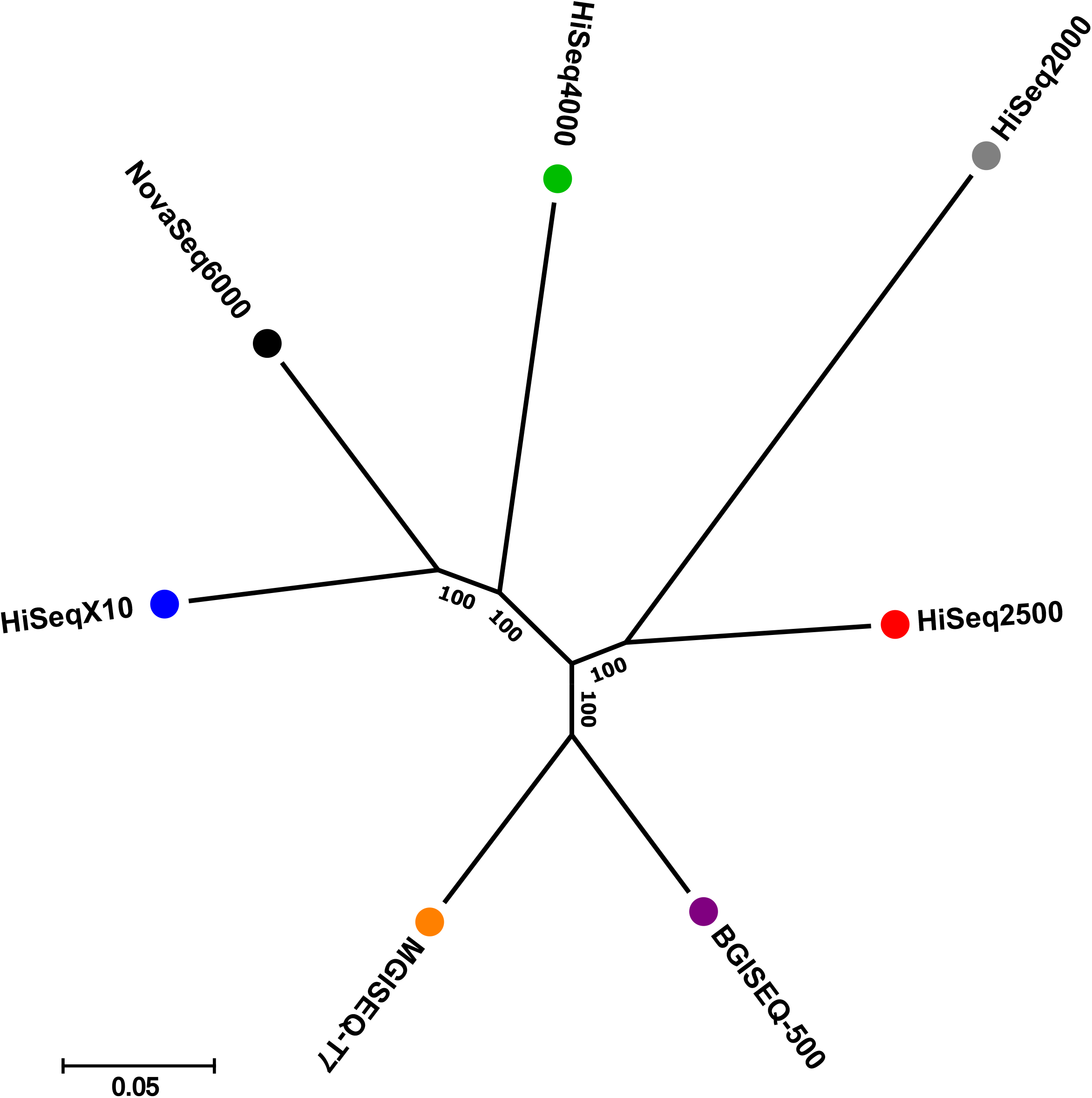
An unrooted tree of seven sequencing platforms showing the similarity of the variant call. Numbers on nodes denote bootstrap values based on 1,000 replicates.

**Table 3.** Variant statistics of Illumina and MGI sequencing platforms.

| Metrics |  | HiSeq2000 | HiSeq2500 | HiSeq4000 | HiSeqX10 | NovaSeq6000 | BGISEQ-500 | MGISEQ-T7 |
| --- | --- | --- | --- | --- | --- | --- | --- | --- |
| Reference homozygous |  | 2,839,356,750 | 2,858,976,629 | 2,855,775,610 | 2,867,632,977 | 2,864,482,967 | 2,855,039,362 | 2,855,211,169 |
| # of no call position |  | 80,242,903 | 60,559,059 | 63,803,179 | 51,824,015 | 54,970,358 | 64,555,525 | 64,401,736 |
| No call rate |  | 2.74% | 2.07% | 2.18% | 1.77% | 1.88% | 2.21% | 2.20% |
| SNVs | Total SNVs | 4,133,415 | 4,197,507 | 4,153,828 | 4,277,851 | 4,283,185 | 4,145,465 | 4,120,925 |
|  | Total SNVs in dbSNP | 4,093,856 | 4,179,089 | 4,128,307 | 4,258,605 | 4,241,561 | 4,125,302 | 4,103,366 |
|  | dbSNP rate | 99.04% | 99.56% | 99.39% | 99.55% | 99.03% | 99.51% | 99.57% |
|  | Singleton | 150,808 | 83,996 | 94,605 | 102,269 | 105,526 | 53,335 | 49,375 |
|  | Singleton in dbSNP | 118,845 | 75,356 | 75,315 | 92,228 | 74,789 | 42,328 | 40,021 |
|  | dbSNP rate for Singleton | 78.81% | 89.71% | 79.61% | 90.18% | 70.87% | 79.36% | 81.06% |
|  | Homozygous | 1,703,636 | 1,697,247 | 1,705,425 | 1,715,123 | 1,720,774 | 1,694,895 | 1,693,653 |
|  | Heterozygous | 2,429,779 | 2,500,260 | 2,448,403 | 2,562,728 | 2,562,411 | 2,450,570 | 2,427,272 |
|  | Het/Hom ratio | 1.43 | 1.47 | 1.44 | 1.49 | 1.49 | 1.45 | 1.43 |
|  | Ti/Tv ratio | 1.91 | 1.9 | 1.9 | 1.87 | 1.84 | 1.91 | 1.91 |
| Indels | Total Indels | 526,451 | 609,968 | 504,179 | 763,447 | 783,294 | 790,152 | 688,728 |
|  | Total Indels in dbSNP | 524,684 | 607,566 | 502,007 | 760,514 | 780,295 | 787,587 | 686,651 |
|  | dbSNP rate | 99.66% | 99.61% | 99.57% | 99.62% | 99.62% | 99.68% | 99.70% |
|  | Singleton | 6,474 | 9,919 | 5,290 | 17,969 | 24,992 | 47,226 | 10,595 |
|  | Singleton in dbSNP | 6,232 | 9,676 | 5,146 | 17,675 | 24,638 | 46,880 | 10,462 |
|  | dbSNP rate for Singleton | 96.26% | 97.55% | 97.28% | 98.36% | 98.58% | 99.27% | 98.74% |

Since it was not possible to conduct standard benchmarking procedures and determine error values for each platform on this study, we compared the variations called by the seven wholegenome sequences with an SNP genotyping chip as the independent platform. Of the total 950,637 comparable positions, more than 99.3% of the genotypes matched the WGS-based genotypes from the seven platforms (Table S9). We found that 4,376 loci in the SNP genotyping were inconsistent across all seven WGS-based genotyping results, suggesting that these loci are probably errors in the SNP genotyping chip. With the exception of HiSeq2000 and HiSeq4000, the other platforms showed a similar concordance rate. This is probably due to the relatively low mapping depth of the HiSeq2000 (28×) and HiSeq4000 (25.8×).

## Discussion

Our benchmark can provide a useful but quite rough estimation of the quality of short-read based whole-genome sequencers. We used the same sample for all the seven sequencers. However, just one human sample cannot justify the variation that may occur in different individuals and DNA molecules and overall sequencing qualities. These are clear limitations, however, as our purpose was to compare two major platforms, still, such a small number of samples can function as an intuitive index for people who consider purchasing expensive sequencers to generate a very large amount of data. Our method of statistical analysis does not allow us to conclude which of the seven sequencing instruments is the most accurate and precise as there is much variation in the sample preparation and sequencer specifications. Nevertheless, overall, the data generated by MGI and Illumina sequencing platforms showed comparable levels of quality, sequencing uniformity, %GC coverage, and concordance rate with SNP genotyping, thus it can be broadly concluded that the MGI platforms can be used for a wide range of research tasks on a par with Illumina platforms at a lower cost.

## Materials and Methods

### Sample and whole-genome sequencing

Genomic DNA used for genotyping and sequencing were extracted from the peripheral blood of Korean male sample donor (KOREF). We constructed sequencing libraries from the KOREF sample for seven different sequencing platforms. We constructed five Illumina sequencing libraries with different insert-sizes (500 bp for HiSeq2000, 400 bp for HiSeq2500 and HiSeq4000, and 450 bp for HiSeqX10 and NovaSeq6000) according to the manufacturer’s protocol (Illumina, San Diego, CA, USA). The Illumina whole-genome sequencing library was sequenced for 90 bp paired-end on HiSeq2000; for 101 bp paired-end on HiSeq2500; for 151 bp paired-end on the HiSeq4000, HiSeqX10 and NovaSeq6000 sequencing platform. We also constructed two MGI sequencing libraries with 300 bp insertsize according to the manufacturer’s protocol [18]. The MGI whole-genome sequencing library was sequenced for 100 bp paired-end on the BGISEQ-500 and MGISEQ-T7 sequencing platform. We conducted genotyping experiments with the KOREF sample using Illumina Infinium Omni1 quad chip according to the manufacturer’s protocols. The Institutional Review Board (IRB) at Ulsan National Institute of Science and Technology approved the study (UNISTIRB-15-19-A).

### Raw data preprocess

We used the FastQC v0.11.8 [6] to assess overall sequencing quality for MGI and Illumina sequencing platforms. PCR duplications (reads were considered duplicates when forward read and reverse read of the two paired-end reads were identical) were detected by the PRINSEQ v0.20.4 (PRINSEQ, RRID:SCR_005454) [19]. The random sequencing error rate was calculated by measuring the occurrence of ‘N’ base at each read position in raw reads. Reads with sequencing adapter contamination were examined according to the manufacturer’s adapter sequences (Illumina sequencing adapter left = *“GATCGGAAGAGCACACGTCTGAACTCCAGTCAC”,* Illumina sequencing adapter right = *“GATCGGAAGAGCGTCGTGTAGGGAAAGAGTGT”,* MGI sequencing adapter left = *“AAGTCGGAGGCCAAGCGGTCTTAGGAAGACAA”,* and MGI sequencing adapter right = *“AAGTCGGATCGTAGCCATGTCGTTCTGTGAGCCAAGGAGTTG”).* We conducted base quality filtration of raw reads using the NGS QC Toolkit v2.3.3 (cutoff read length for high quality 70; cutoff quality score, 20) (NGS QC Toolkit, RRID:SCR_005461) [20]. We used clean reads after removing low-quality reads and adapter containing reads for the mapping step.

### Mapping, variant calling, and coverage calculation

After the filtering step, clean reads were aligned to the human reference genome (GRCh38) using BWA-MEM v0.7.12, and duplicates marked with Picard v2.6.0 (Picard, RRID:SCR_006525) [21]. Realignment and base score recalibration of the bam file was processed by GATK v3.3. Single nucleotide variants, short insertions, and deletions were called with the GATK (Unifiedgenotyper, options --output_mode EMIT_ALL_SITES -- genotype_likelihoods_model BOTH). The resulting variants were annotated with the dbSNP (v153) database [22]. Coverage was calculated for each nucleotide using SAMtools v1.9 (SAMTOOLS, RRID:SCR_002105) [23]. We defined a specific covered region based on the 100 bp non-overlapping windows by calculating the average depth of the windows. We used more than five times the difference with an average depth in each window between MGI and Illumina platforms. GC coverage for raw reads and genome was calculated by the average %GC of the 100bp non-overlapping windows.

### Variants comparison and concordance rate with SNP genotyping

The chromosome position and genotype of each variant called from each sequencing platform was used to identify the relationship between seven sequencing platforms. We compared 1,034,447 loci found on one or more platforms for locations where genotypes were determined on all platforms. An unrooted tree was generated using FastTree v2.1.10 (FastTree, RRID:SCR_015501) [24] with the generalized time-reversible (GTR) model. For calculating the concordance rate between SNP genotyping and WGS-based genotype, the coordinates of SNP genotyping data were converted to GRCh38 assembly using the UCSC LiftOver tool [25]. We removed unmapped positions and indel markers and used only markers that were present on the autosomal chromosomes.

## Availability of Supporting Data and Materials

All sequences generated in this study, including the HiSeq2000, HiSeq2500, HiSeq4000, HiSeqX10, NovaSeq6000, BGISEQ-500, and MGISEQ-T7 sequencing reads, were deposited in the NCBI Sequence Read Archive database under BioProject PRJNA600063. All the data will be hosted and distributed from http://biosequencer.org.

## Additional Files

Additional file 1: **Figure S1**. Distribution of nucleotide quality across seven sequencing platforms. **Figure S2**. Base quality filtration statistics of seven sequencing platforms. **Figure S3**. Random error ratio in seven sequencing platforms. **Figure S4**. Insert-size distributions of seven sequencing platforms. **Figure S5**. The coverage distribution of two MGI and five Illumina platforms. **Figure S6**. Depth distribution of chromosome 8. **Figure S7**. Depth distribution of chromosome 12. **Figure S8**. Depth distribution of chromosome 18. **Figure S9**. Depth distribution of chromosome 20. **Figure S10**. GC distribution of platform-specific covered region. **Figure S11**. The GC composition distribution of the human genome and sequencing reads. **Table S1**. Base quality summary. **Table S2**. Duplicate reads, random error base, and adapter read rate. **Table S3**. Statistics of clean reads for seven sequencing platforms. **Table S4**. Statistics of platform-specific covered regions. **Table S5**. The number of shared SNVs in seven platforms. **Table S6**. The number of SNVs that were singleton or not found in a specific platform. **Table S7**. Genotype concordance rate among seven sequencing platforms. **Table S8**. Genotype comparison between SNP genotyping and WGS.

## List of abbreviations

PE: paired-end;
WGS: whole-genome sequencing;
BWA: burrows-wheeler aligner;
SNVs: single nucleotide variants;
indels: insertions and deletions;
Ti/Tv: transition/transversion;
GATK: Genome Analysis ToolKit;

## Authors’ contributions

J.B. supervised and coordinated the project. J.B. and Y.S.C. conceived and designed the experiments. H.M.K., S.J., O.C., J.H.J., H.Y.L., and Y.Y. conducted the bioinformatics data processing and analyses. H.M.K., S.J., D.M.B., and J.B. wrote and revised the manuscript. A.S. and H.S.K. reviewed and edited the manuscript. All authors read and approved the final manuscript.

## Acknowledgments

We thank the Korea Institute of Science and Technology Information (KISTI) provided us the Korea Research Environment Open NETwork (KREONET). We thank Jaesu Bhak for editing.

