## Supplementary Figures and Tables for "Comparative analysis of seven short-reads sequencing platforms using the Korean Reference Genome: MGI and Illumina sequencing benchmark for whole-genome sequencing"

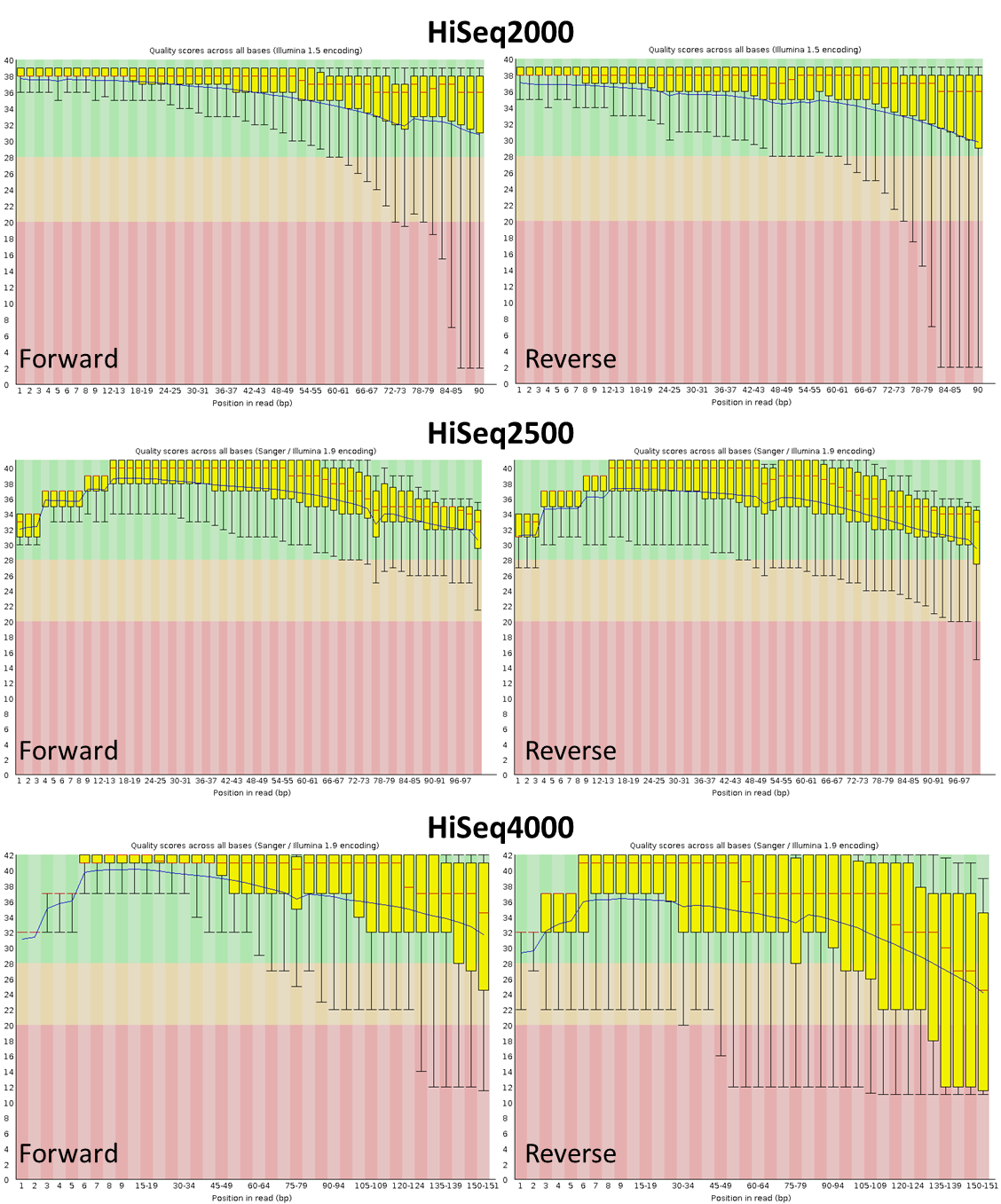

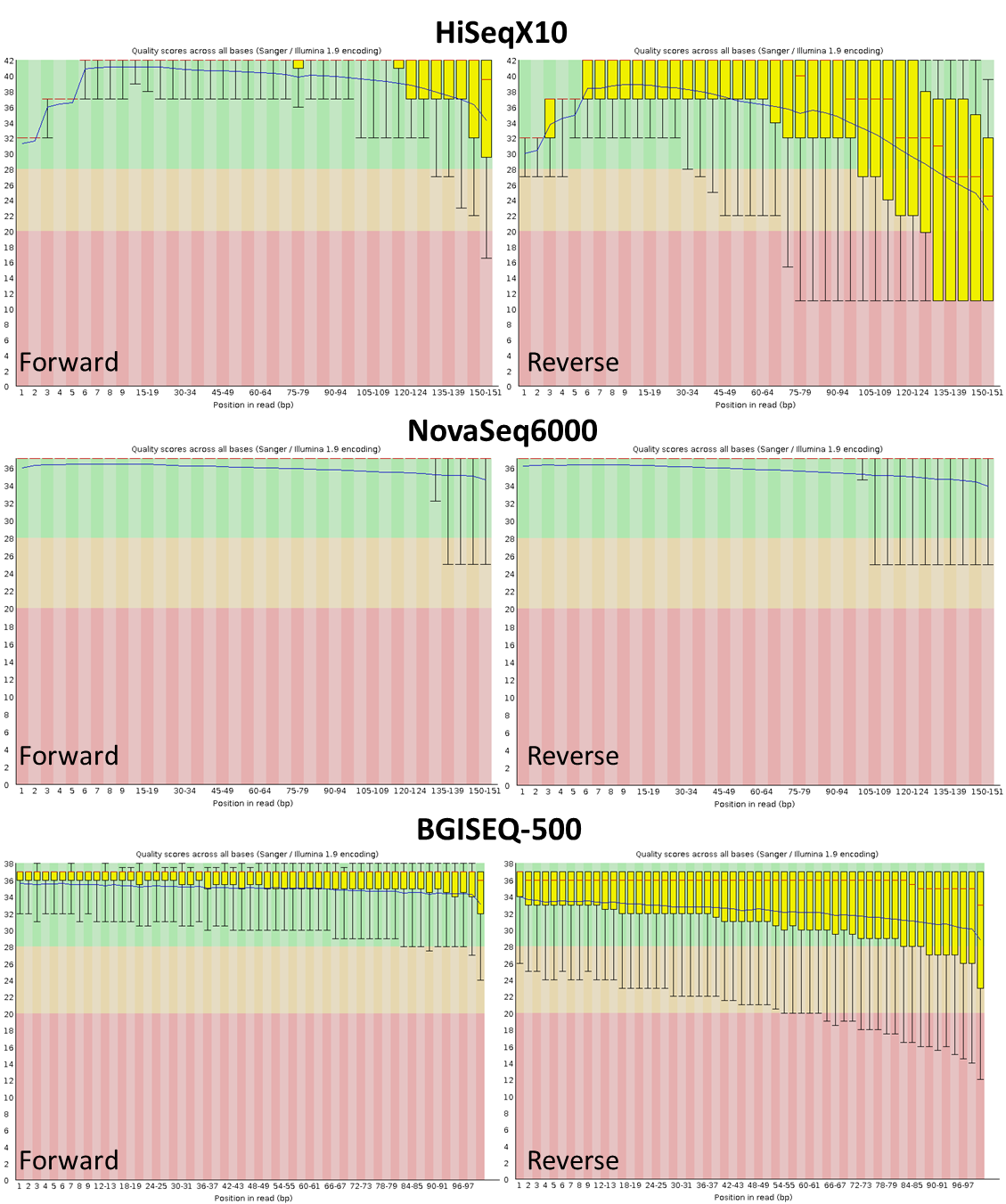

**Figure S1. Distribution of nucleotide quality across seven sequencing platforms.**
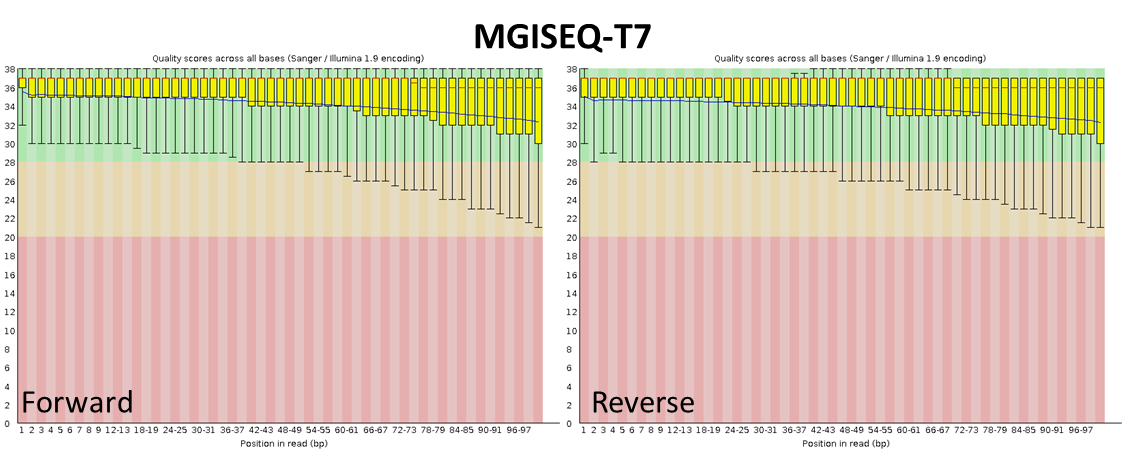

**Figure S2. Base quality filtration statistics of seven sequencing platforms**
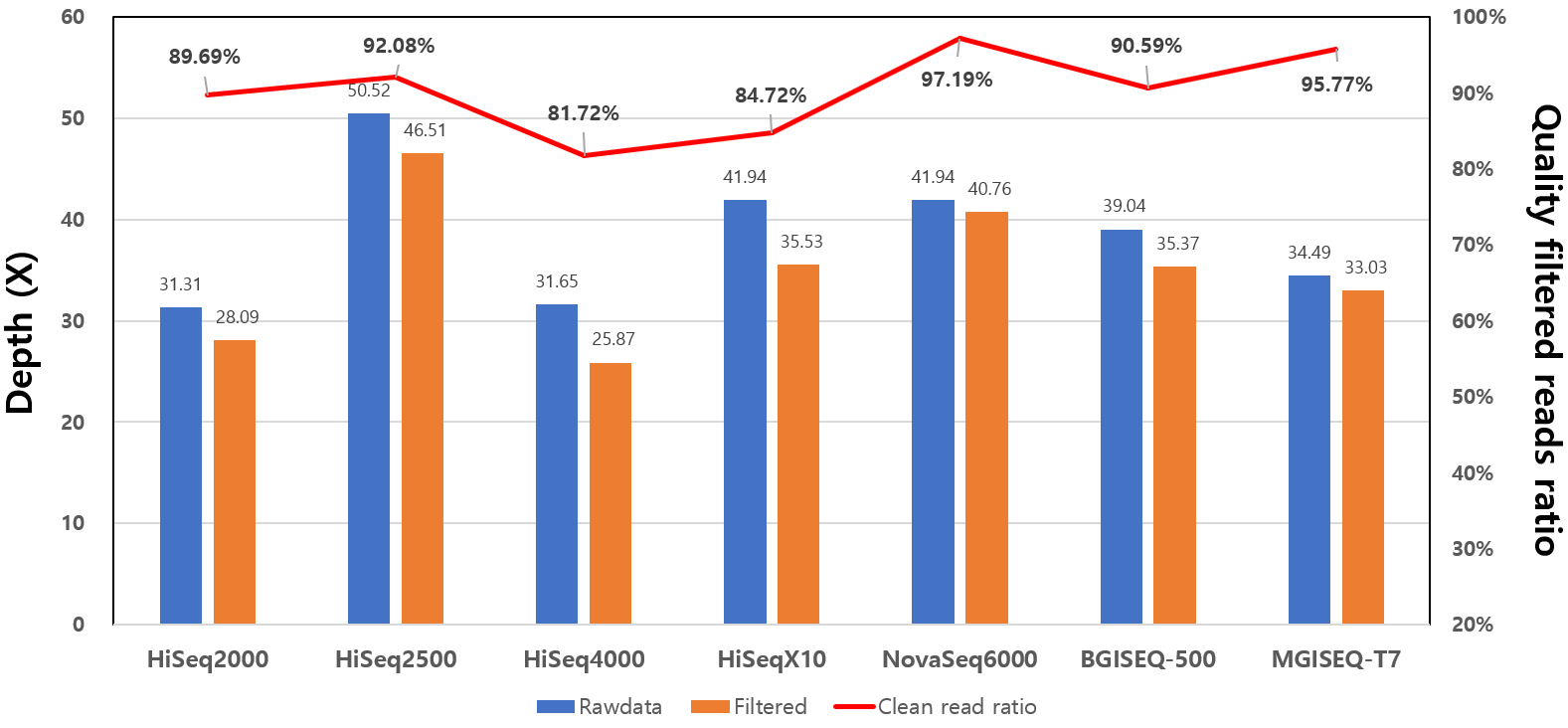

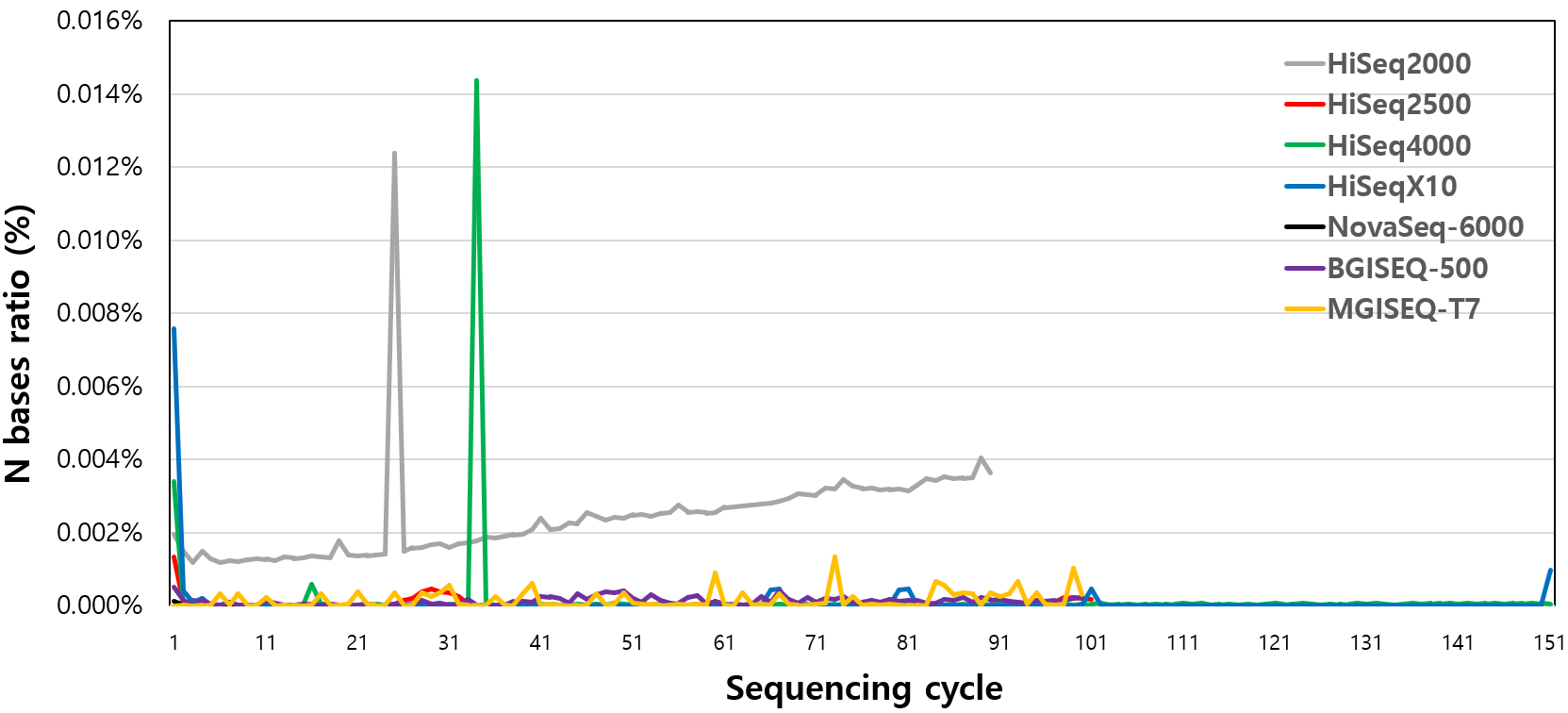

**Figure S3. Random error ratio in seven sequencing platforms.**

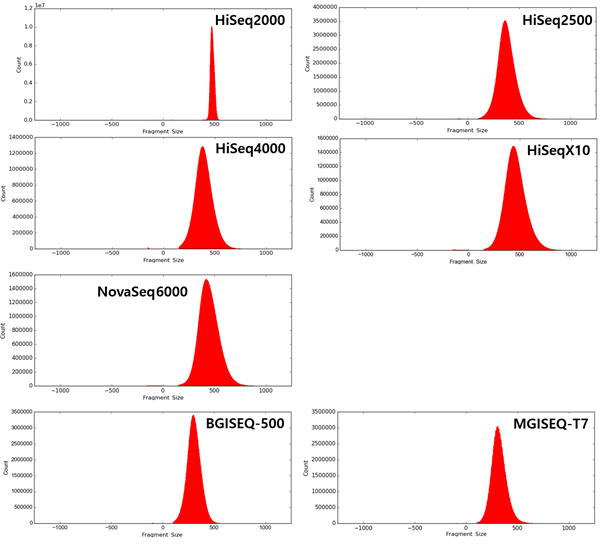

**Figure S4. Insert-size distributions of seven sequencing platforms**

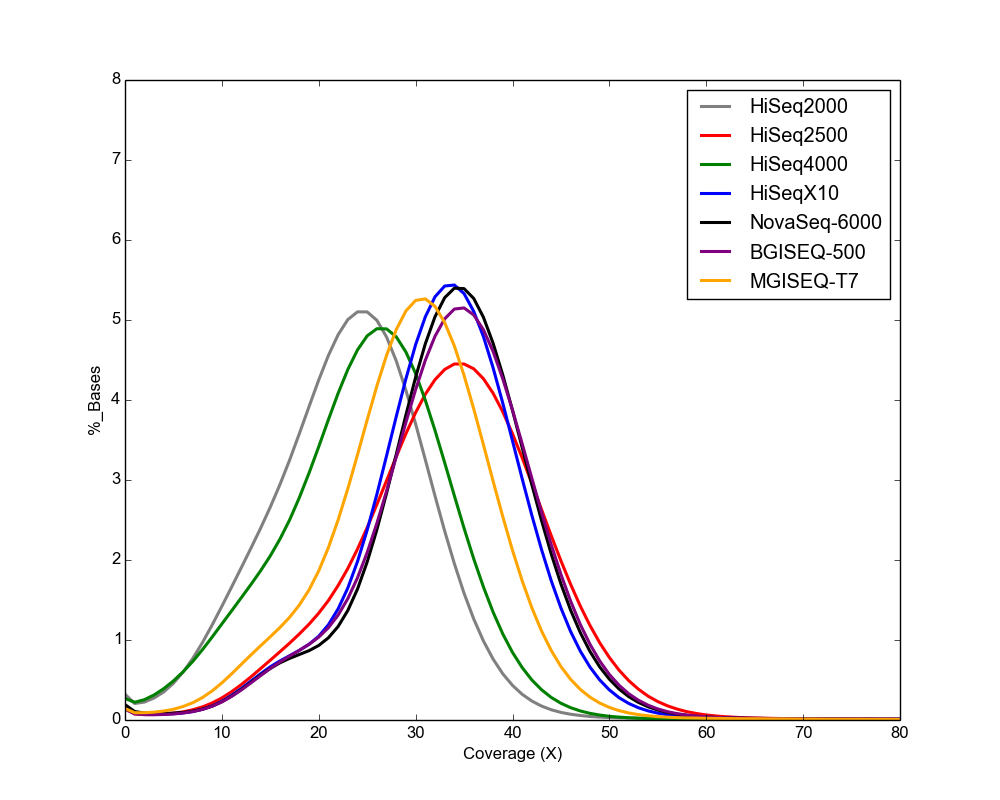

**Figure S5. The coverage distribution of two MGI and five Illumina platforms.**

**Figure S6. Depth distribution of chromosome 8**
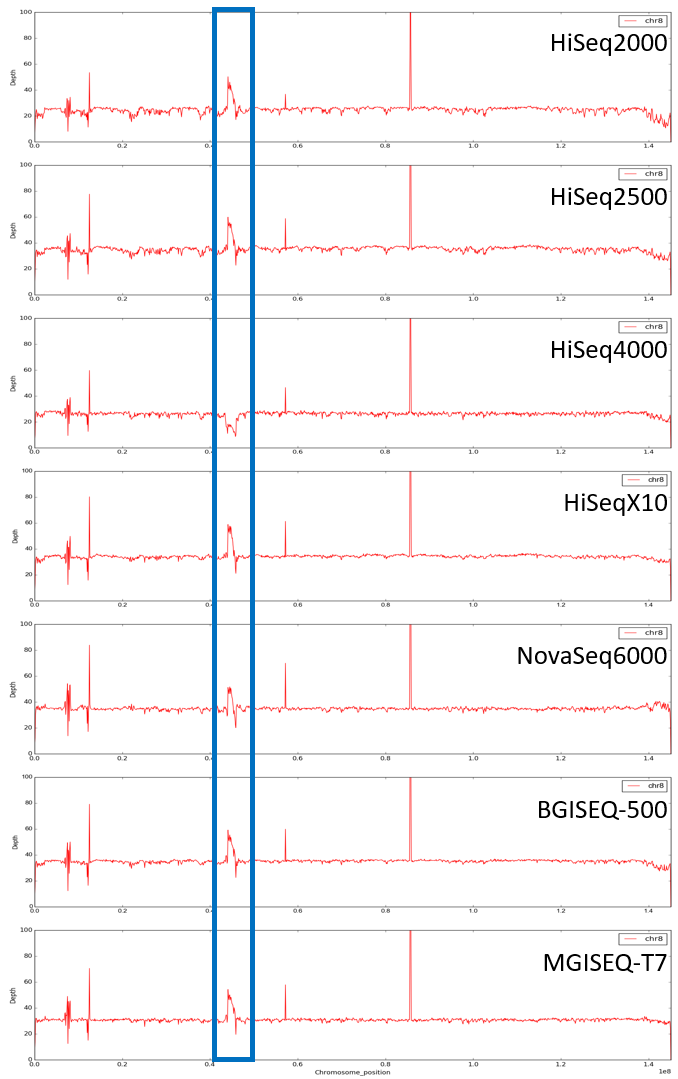

**Figure S7. Depth distribution of chromosome 12**
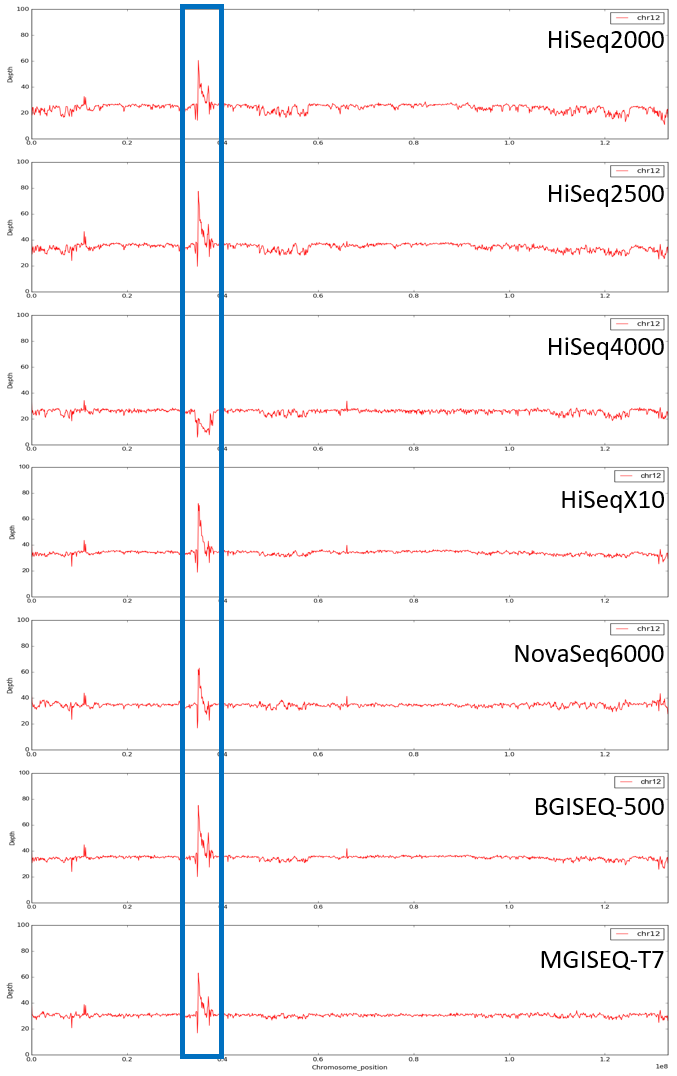

**Figure S8. Depth distribution of chromosome 18**
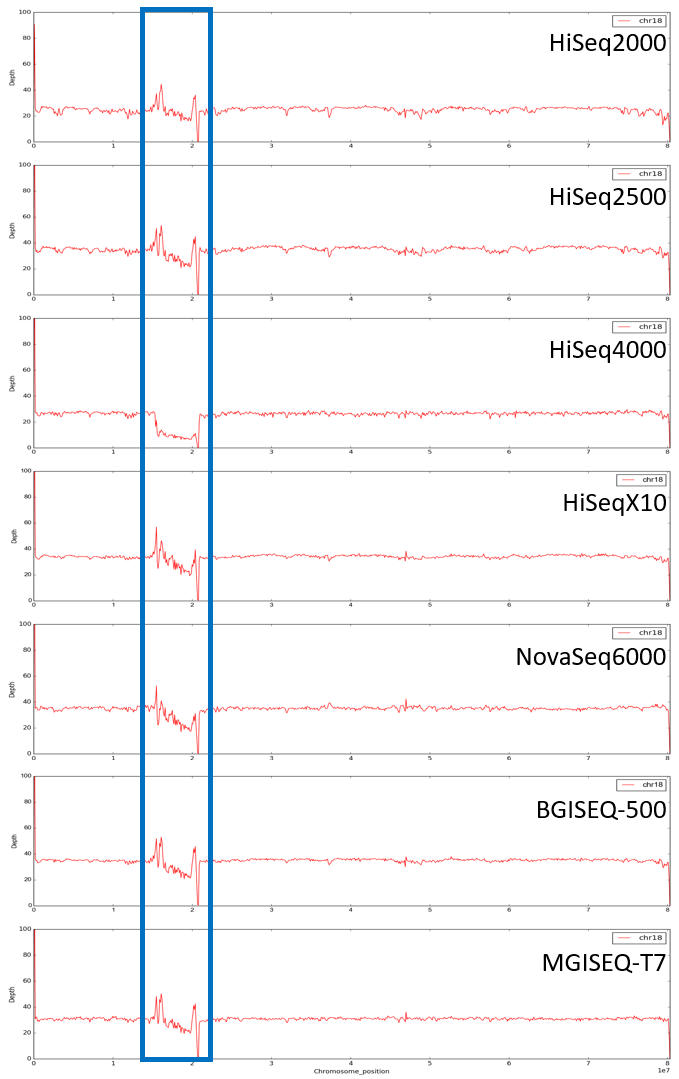

**Figure S9. Depth distribution of chromosome 20**
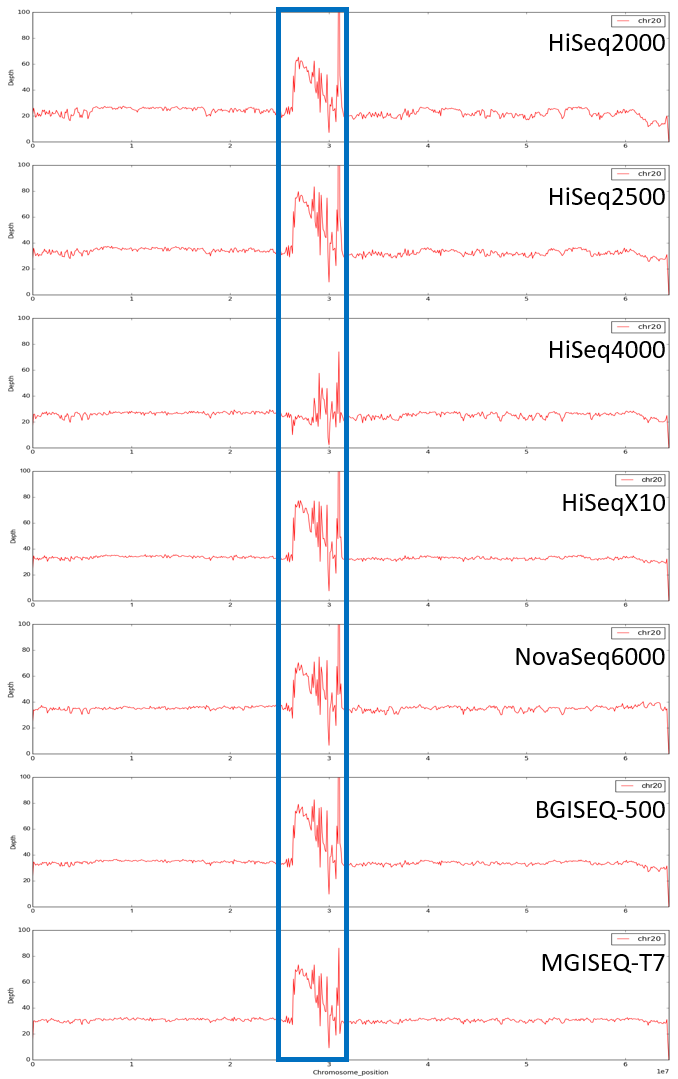

**Figure S10. GC distribution of the platform-specific covered region**
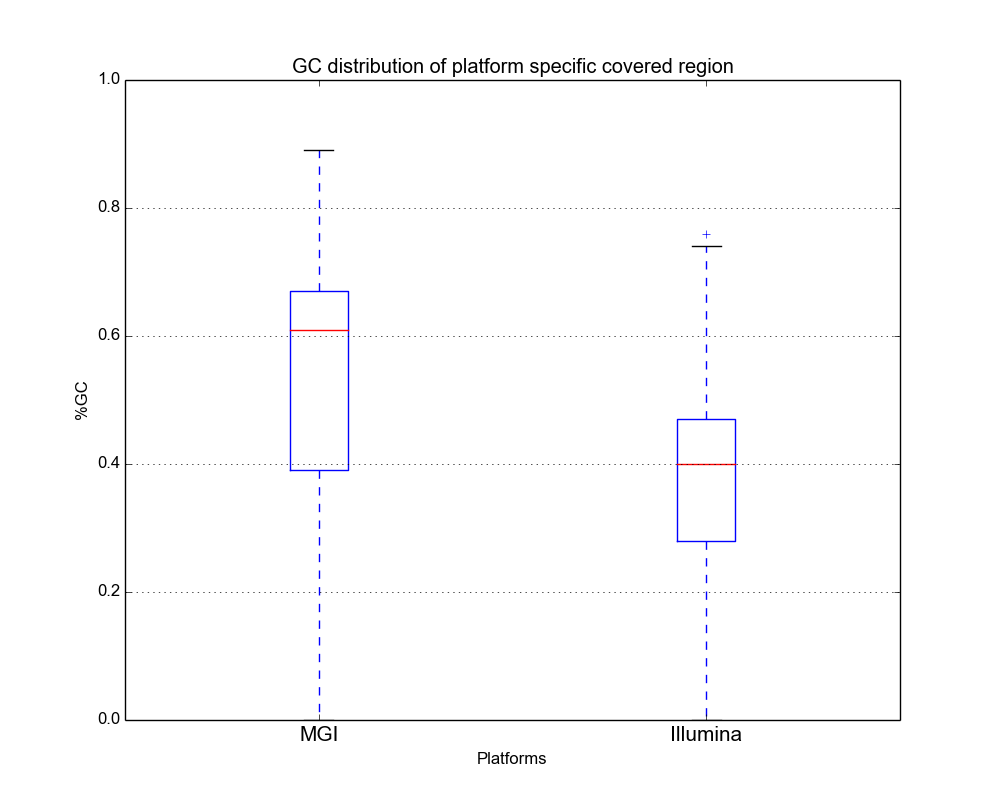

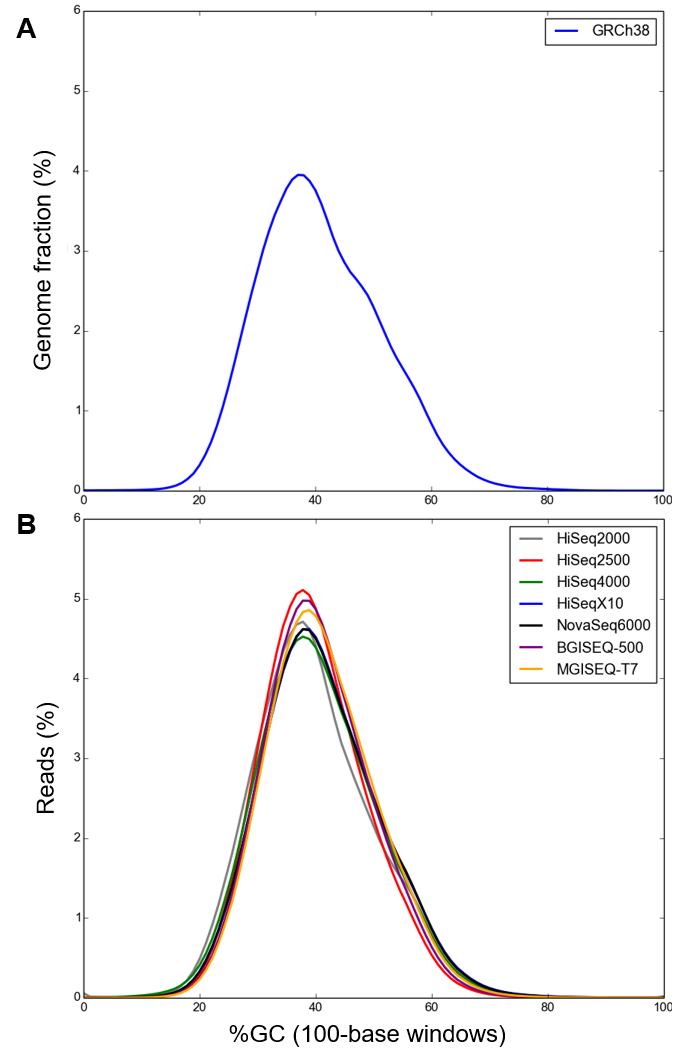

**Figure S11. The GC composition distribution of the human genome and sequencing reads.** (A) The GC composition distribution of the human genome (GRCh38). (B) The GC composition distribution of sequencing reads from seven sequencing platforms.

**Supplementary Tables**

**Table S1. Base quality summary**

| Platforms | Total number  of raw reads | Raw reads >Q20 bases | Raw reads >Q30 bases | Quality filtered read | % of clean reads |
| --- | --- | --- | --- | --- | --- |
| HiSeq2000 | 1,043,812,460 | 94.26% | 88.13% | 936,199,954 | 89.69% |
| HiSeq2500 | 1,500,482,864 | 93.92% | 83.35% | 1,381,620,954 | 92.08% |
| HiSeq4000 | 628,832,102 | 81.01% | 61.10% | 513,909,986 | 81.72% |
| HiSeqX10 | 833,299,268 | 82.73% | 64.72% | 705,987,474 | 84.72% |
| NovaSeq6000 | 833,299,268 | 96.94% | 92.31% | 809,888,988 | 97.19% |
| BGISEQ-500 | 1,171,205,242 | 93.45% | 81.93% | 1,061,036,526 | 90.59% |
| MGISEQ-T7 | 1,034,757,668 | 94.97% | 83.55% | 991,023,340 | 95.77% |

**Table S2. Duplicate reads, random error base, and adapter read rate**

|  | Duplicate  reads rate | Random base  error rate | Adapter  contamination rate* |
| --- | --- | --- | --- |
| HiSeq2000 | 8.71% | 0.22% | 0.03% |
| HiSeq2500 | 1.66% | 0.00% | 0.03% |
| HiSeq4000 | 0.67% | 0.02% | 0.24% |
| HiSeqX10 | 1.21% | 0.01% | 0.23% |
| NovaSeq6000 | 1.70% | 0.00% | 0.21% |
| BGISEQ-500 | 0.68% | 0.01% | 0.02% |
| MGISEQ-T7 | 3.04% | 0.02% | 0.00% |

* Adapter reads of MGISEQ-T7 were already filtered out at the instrument.

**Table S3. The putatively erroneous *K*-mers (≤ 3 *K*-mer depth) in seven sequencing platforms.**

| *K*-mer  depth | Proportion of putatively erroneous *K*-mer | | | | | | |
| --- | --- | --- | --- | --- | --- | --- | --- |
|  | HiSeq2000 | HiSeq2500 | HiSeq4000 | HiSeqX10 | NovaSeq6000 | BGISEQ-500 | MGISEQ-T7 |
| 1 | 3.83% | 5.32% | 12.38% | 10.98% | 3.54% | 7.12% | 5.84% |
| 2 | 0.39% | 0.38% | 1.21% | 0.85% | 0.28% | 0.48% | 0.45% |
| 3 | 0.16% | 0.12% | 0.32% | 0.22% | 0.09% | 0.12% | 0.10% |
| Sum | 4.38% | 5.82% | 13.91% | 12.05% | 3.91% | 7.72% | 6.39% |

**Table S4. Statistics of clean reads for seven sequencing platforms.**

| Platforms | Raw read  depth (×) | Clean read  depth (×) | Down-sampled  depth (×) |
| --- | --- | --- | --- |
| HiSeq2000 | 31.31 | 28.08 | - |
| HiSeq2500 | 50.52 | 46.50 | 35.35 |
| HiSeq4000 | 31.65 | 25.82 | - |
| HiSeqX10 | 41.94 | 35.53 | - |
| NovaSeq6000 | 41.94 | 40.70 | 35.54 |
| BGISEQ-500 | 39.04 | 35.36 | - |
| MGISEQ-T7 | 34.49 | 33.03 | - |

*The depths shown in table were calculated based on 3 Gb.

**Table S5. Statistics of platform-specific covered regions**

|  | MGI platforms (2) | Illumina platforms (5) |
| --- | --- | --- |
| Specific covered region counts* | 1,516 | 2,264 |
| Specific covered region length (bp) | 151,600 | 226,400 |
| # of genes in specific covered region | 243 | 717 |
| # of CDS affected by specific covered region | 8 | 0 |

*Specific covered region was calculated by comparing the average depth of MGI and Illumina platforms based on 100 bp nonoverlapping windows.

**Table S6. The number of shared SNVs in seven platforms**

| **Shared Platforms** | **Count** |
| --- | --- |
| HiSeq2000_HiSeq2500_HiSeq4000_HiSeqX10_NovaSeq6000_BGISEQ-500_MGISEQ-T7 | 3,744,554 |
| HiSeq2500_HiSeq4000_HiSeqX10_NovaSeq6000_BGISEQ-500_MGISEQ-T7 | 79,982 |
| HiSeq2000_HiSeq2500_HiSeqX10_NovaSeq6000_BGISEQ-500_MGISEQ-T7 | 35,322 |
| HiSeqX10_NovaSeq6000 | 33,902 |
| HiSeq4000_HiSeqX10_NovaSeq6000 | 28,485 |
| HiSeq2000_HiSeq2500_HiSeq4000_HiSeqX10_NovaSeq6000_BGISEQ-500 | 16,328 |
| HiSeq2000_HiSeq2500 | 15,936 |
| BGISEQ-500_MGISEQ-T7 | 15,670 |
| HiSeq4000_HiSeqX10 | 12,980 |
| HiSeq4000_NovaSeq6000 | 12,166 |
| HiSeq2500_BGISEQ-500_MGISEQ-T7 | 12,014 |
| HiSeq2500_HiSeqX10_NovaSeq6000_BGISEQ-500_MGISEQ-T7 | 11,739 |
| HiSeq4000_HiSeqX10_NovaSeq6000_BGISEQ-500_MGISEQ-T7 | 11,314 |
| HiSeq2000_HiSeq2500_HiSeq4000_HiSeqX10_NovaSeq6000_MGISEQ-T7 | 10,595 |
| HiSeq2000_HiSeq2500_HiSeq4000_HiSeqX10_NovaSeq6000 | 9,325 |
| HiSeq2500_BGISEQ-500 | 8,790 |
| HiSeq2500_HiSeq4000_HiSeqX10_NovaSeq6000_BGISEQ-500 | 8,501 |
| HiSeq2500_HiSeq4000_HiSeqX10_NovaSeq6000 | 8,106 |
| HiSeq2000_HiSeqX10 | 8,104 |
| HiSeq2000_NovaSeq6000 | 7,761 |
| HiSeq2500_MGISEQ-T7 | 7,692 |
| HiSeq2500_HiSeqX10 | 7,624 |
| HiSeq2000_HiSeq4000_HiSeqX10_NovaSeq6000_BGISEQ-500_MGISEQ-T7 | 7,569 |
| HiSeq2500_HiSeq4000_HiSeqX10_NovaSeq6000_MGISEQ-T7 | 6,986 |
| HiSeq2000_HiSeqX10_NovaSeq6000 | 6,794 |
| HiSeq2000_HiSeq2500_BGISEQ-500_MGISEQ-T7 | 6,617 |
| HiSeq4000_HiSeqX10_NovaSeq6000_BGISEQ-500 | 6,612 |
| HiSeq2000_HiSeq4000_HiSeqX10_NovaSeq6000 | 6,468 |
| HiSeq2500_HiSeqX10_NovaSeq6000 | 6,326 |
| HiSeq2000_HiSeq2500_HiSeqX10_NovaSeq6000 | 6,159 |
| HiSeq2500_NovaSeq6000 | 6,056 |
| HiSeq2000_BGISEQ-500 | 5,517 |
| HiSeqX10_NovaSeq6000_BGISEQ-500_MGISEQ-T7 | 5,474 |
| NovaSeq6000_BGISEQ-500 | 5,410 |
| HiSeq2000_MGISEQ-T7 | 5,310 |
| HiSeq2000_HiSeq2500_HiSeqX10_NovaSeq6000_BGISEQ-500 | 5,302 |
| HiSeq4000_HiSeqX10_NovaSeq6000_MGISEQ-T7 | 5,241 |
| HiSeqX10_BGISEQ-500 | 4,866 |
| HiSeqX10_NovaSeq6000_BGISEQ-500 | 4,752 |
| HiSeq2500_NovaSeq6000_BGISEQ-500_MGISEQ-T7 | 4,627 |
| NovaSeq6000_MGISEQ-T7 | 4,625 |
| HiSeq2500_HiSeq4000 | 4,525 |
| HiSeq2000_HiSeq2500_HiSeq4000_NovaSeq6000_BGISEQ-500_MGISEQ-T7 | 4,484 |
| HiSeq2000_HiSeq2500_HiSeq4000_HiSeqX10_BGISEQ-500_MGISEQ-T7 | 4,237 |
| HiSeq2500_HiSeq4000_NovaSeq6000_BGISEQ-500_MGISEQ-T7 | 4,138 |
| HiSeq2000_HiSeq4000 | 4,128 |
| HiSeq2000_HiSeq2500_HiSeqX10 | 4,056 |
| HiSeqX10_MGISEQ-T7 | 4,053 |
| HiSeq2500_HiSeqX10_NovaSeq6000_BGISEQ-500 | 3,874 |
| HiSeq2000_HiSeq2500_BGISEQ-500 | 3,854 |
| NovaSeq6000_BGISEQ-500_MGISEQ-T7 | 3,843 |
| HiSeq2000_HiSeq2500_NovaSeq6000_BGISEQ-500_MGISEQ-T7 | 3,558 |
| HiSeqX10_NovaSeq6000_MGISEQ-T7 | 3,555 |
| HiSeq2000_HiSeq2500_HiSeqX10_NovaSeq6000_MGISEQ-T7 | 3,491 |
| HiSeq2500_HiSeqX10_BGISEQ-500_MGISEQ-T7 | 3,452 |
| HiSeq2000_HiSeq2500_MGISEQ-T7 | 3,358 |
| HiSeq2000_HiSeq2500_HiSeqX10_BGISEQ-500_MGISEQ-T7 | 3,337 |
| HiSeq2500_HiSeq4000_HiSeqX10_BGISEQ-500_MGISEQ-T7 | 3,325 |
| HiSeq4000_BGISEQ-500 | 3,312 |
| HiSeq2000_HiSeq2500_NovaSeq6000 | 3,244 |
| HiSeq2000_HiSeqX10_NovaSeq6000_BGISEQ-500_MGISEQ-T7 | 3,124 |
| HiSeqX10_BGISEQ-500_MGISEQ-T7 | 2,992 |
| HiSeq4000_MGISEQ-T7 | 2,950 |
| HiSeq2000_BGISEQ-500_MGISEQ-T7 | 2,932 |
| HiSeq2500_HiSeqX10_NovaSeq6000_MGISEQ-T7 | 2,931 |
| HiSeq2000_HiSeq4000_HiSeqX10_NovaSeq6000_BGISEQ-500 | 2,705 |
| HiSeq2500_HiSeq4000_BGISEQ-500_MGISEQ-T7 | 2,516 |
| HiSeq4000_BGISEQ-500_MGISEQ-T7 | 2,484 |
| HiSeq2500_HiSeq4000_HiSeqX10 | 2,384 |
| HiSeq2500_NovaSeq6000_BGISEQ-500 | 2,376 |
| HiSeq2000_HiSeq4000_HiSeqX10_NovaSeq6000_MGISEQ-T7 | 2,313 |
| HiSeq4000_NovaSeq6000_BGISEQ-500_MGISEQ-T7 | 2,312 |
| HiSeq2500_HiSeqX10_BGISEQ-500 | 2,205 |
| HiSeq2500_NovaSeq6000_MGISEQ-T7 | 2,073 |
| HiSeq2500_HiSeq4000_NovaSeq6000 | 2,072 |
| HiSeq4000_NovaSeq6000_BGISEQ-500 | 2,033 |
| HiSeq2000_HiSeq2500_HiSeq4000 | 1,974 |
| HiSeq2000_HiSeq4000_HiSeqX10 | 1,926 |
| HiSeq2000_HiSeq2500_HiSeq4000_BGISEQ-500_MGISEQ-T7 | 1,897 |
| HiSeq4000_HiSeqX10_BGISEQ-500 | 1,897 |
| HiSeq2000_HiSeq4000_NovaSeq6000 | 1,884 |
| HiSeq2000_HiSeqX10_NovaSeq6000_BGISEQ-500 | 1,865 |
| HiSeq4000_HiSeqX10_BGISEQ-500_MGISEQ-T7 | 1,865 |
| HiSeq2500_HiSeqX10_MGISEQ-T7 | 1,839 |
| HiSeq4000_NovaSeq6000_MGISEQ-T7 | 1,728 |
| HiSeq2000_HiSeq2500_HiSeqX10_BGISEQ-500 | 1,669 |
| HiSeq2000_HiSeq2500_HiSeq4000_HiSeqX10 | 1,637 |
| HiSeq2000_HiSeq2500_NovaSeq6000_BGISEQ-500 | 1,627 |
| HiSeq4000_HiSeqX10_MGISEQ-T7 | 1,591 |
| HiSeq2000_HiSeqX10_NovaSeq6000_MGISEQ-T7 | 1,466 |
| HiSeq2000_NovaSeq6000_BGISEQ-500 | 1,425 |
| HiSeq2000_HiSeq2500_NovaSeq6000_MGISEQ-T7 | 1,397 |
| HiSeq2000_HiSeq2500_HiSeqX10_MGISEQ-T7 | 1,392 |
| HiSeq2000_HiSeq2500_HiSeq4000_NovaSeq6000 | 1,388 |
| HiSeq2500_HiSeq4000_NovaSeq6000_BGISEQ-500 | 1,367 |
| HiSeq2000_HiSeq2500_HiSeq4000_NovaSeq6000_BGISEQ-500 | 1,280 |
| HiSeq2000_HiSeq2500_HiSeq4000_HiSeqX10_BGISEQ-500 | 1,274 |
| HiSeq2500_HiSeq4000_HiSeqX10_BGISEQ-500 | 1,261 |
| HiSeq2500_HiSeq4000_BGISEQ-500 | 1,258 |
| HiSeq2500_HiSeq4000_HiSeqX10_MGISEQ-T7 | 1,213 |
| HiSeq2000_HiSeqX10_BGISEQ-500 | 1,210 |
| HiSeq2000_NovaSeq6000_MGISEQ-T7 | 1,200 |
| HiSeq2000_NovaSeq6000_BGISEQ-500_MGISEQ-T7 | 1,193 |
| HiSeq2500_HiSeq4000_NovaSeq6000_MGISEQ-T7 | 1,173 |
| HiSeq2500_HiSeq4000_MGISEQ-T7 | 1,099 |
| HiSeq2000_HiSeqX10_MGISEQ-T7 | 1,033 |
| HiSeq2000_HiSeqX10_BGISEQ-500_MGISEQ-T7 | 1,007 |
| HiSeq2000_HiSeq2500_HiSeq4000_HiSeqX10_MGISEQ-T7 | 1,000 |
| HiSeq2000_HiSeq2500_HiSeq4000_NovaSeq6000_MGISEQ-T7 | 966 |
| HiSeq2000_HiSeq2500_HiSeq4000_BGISEQ-500 | 814 |
| HiSeq2000_HiSeq4000_NovaSeq6000_BGISEQ-500_MGISEQ-T7 | 756 |
| HiSeq2000_HiSeq4000_HiSeqX10_BGISEQ-500_MGISEQ-T7 | 732 |
| HiSeq2000_HiSeq4000_BGISEQ-500 | 729 |
| HiSeq2000_HiSeq2500_HiSeq4000_MGISEQ-T7 | 725 |
| HiSeq2000_HiSeq4000_MGISEQ-T7 | 660 |
| HiSeq2000_HiSeq4000_NovaSeq6000_BGISEQ-500 | 606 |
| HiSeq2000_HiSeq4000_BGISEQ-500_MGISEQ-T7 | 591 |
| HiSeq2000_HiSeq4000_HiSeqX10_BGISEQ-500 | 564 |
| HiSeq2000_HiSeq4000_NovaSeq6000_MGISEQ-T7 | 541 |
| HiSeq2000_HiSeq4000_HiSeqX10_MGISEQ-T7 | 507 |

**Table S7. The number of SNVs that were singleton or not found in a specific platform.**

| Detection of SNVs | | # of SNVs |
| --- | --- | --- |
| Found | Not found |  |
| HiSeq2000 (singleton) | HiSeq2500_HiSeq4000_HiSeqX10_NovaSeq6000_BGISEQ-500_MGISEQ-T7 | 150,808 |
| NovaSeq6000 (singleton) | HiSeq2000_HiSeq2500_HiSeq4000_HiSeqX10_BGISEQ-500_MGISEQ-T7 | 105,526 |
| HiSeqX10 (singleton) | HiSeq2000_HiSeq2500_HiSeq4000_NovaSeq6000_BGISEQ-500_MGISEQ-T7 | 102,269 |
| HiSeq4000 (singleton) | HiSeq2000_HiSeq2500_HiSeqX10_NovaSeq6000_BGISEQ-500_MGISEQ-T7 | 94,605 |
| HiSeq2500 (singleton) | HiSeq2000_HiSeq4000_HiSeqX10_NovaSeq6000_BGISEQ-500_MGISEQ-T7 | 83,996 |
| BGISEQ-500 (singleton) | HiSeq2000_HiSeq2500_HiSeq4000_HiSeqX10_NovaSeq6000_MGISEQ-T7 | 53,335 |
| MGISEQ-T7 (singleton) | HiSeq2000_HiSeq2500_HiSeq4000_HiSeqX10_NovaSeq6000_BGISEQ-500 | 49,375 |
| HiSeq2500_HiSeq4000_HiSeqX10_NovaSeq6000_BGISEQ-500_MGISEQ-T7 | HiSeq2000 | 79,982 |
| HiSeq2000_HiSeq2500_HiSeqX10_NovaSeq6000_BGISEQ-500_MGISEQ-T7 | HiSeq4000 | 35,322 |
| HiSeq2000_HiSeq2500_HiSeq4000_HiSeqX10_NovaSeq6000_BGISEQ-500 | MGISEQ-T7 | 16,328 |
| HiSeq2000_HiSeq2500_HiSeq4000_HiSeqX10_NovaSeq6000_MGISEQ-T7 | BGISEQ-500 | 10,595 |
| HiSeq2000_HiSeq4000_HiSeqX10_NovaSeq6000_BGISEQ-500_MGISEQ-T7 | HiSeq2500 | 7,569 |
| HiSeq2000_HiSeq2500_HiSeq4000_NovaSeq6000_BGISEQ-500_MGISEQ-T7 | HiSeqX10 | 4,484 |
| HiSeq2000_HiSeq2500_HiSeq4000_HiSeqX10_BGISEQ-500_MGISEQ-T7 | NovaSeq6000 | 4,237 |

**Table S8. Genotype concordance rate among seven sequencing platforms**

| Platforms | HiSeq2500 | HiSeq4000 | HiSeqX10 | NovaSeq6000 | BGISEQ-500 | MGISEQ-T7 |
| --- | --- | --- | --- | --- | --- | --- |
| HiSeq2000 | 56.85% | 48.08% | 51.46% | 50.95% | 56.41% | 56.27% |
| HiSeq2500 | - | 59.08% | 63.23% | 62.68% | 70.38% | 69.74% |
| HiSeq4000 | - | - | 63.63% | 63.18% | 62.19% | 62.01% |
| HiSeqX10 | - | - | - | 70.68% | 65.31% | 64.49% |
| NovaSeq6000 | - | - | - | - | 65.53% | 64.64% |
| BGISEQ-500 | - | - | - | - | - | 76.61% |

**Table S9. Genotype comparison between SNP genotyping and WGS**

| Platforms | Total comparable  positions in  SNP genotyping | Match  count | Mismatch  Count* | Genotype concordance rate |
| --- | --- | --- | --- | --- |
| HiSeq2000 | 950,637 | 944,618 | 6,019 | 99.37% |
| HiSeq2500 | 950,637 | 946,006 | 4,631 | 99.51% |
| HiSeq4000 | 950,637 | 945,432 | 5,205 | 99.45% |
| HiSeqX10 | 950,637 | 946,056 | 4,581 | 99.52% |
| NovaSeq6000 | 950,637 | 946,060 | 4,577 | 99.52% |
| BGISEQ-500 | 950,637 | 946,045 | 4,592 | 99.52% |
| MGISEQ-T7 | 950,637 | 946,011 | 4,626 | 99.51% |

* 4,376 loci were mismatched in all seven platforms.
